# Fine stratification of microbial communities through a metagenomic profile of the photic zone

**DOI:** 10.1101/134635

**Authors:** Jose M. Haro-Moreno, Mario López-Pérez, José R. de la Torre, Antonio Picazo, Antonio Camacho, Francisco Rodríguez-Valera

**Affiliations:** Evolutionary Genomics Group, División de Microbiología, Universidad Miguel Hernández, Apartado 18, San Juan 03550, Alicante, Spain; Department of Biology, San Francisco State University, San Francisco, CA 94132, USA; Cavanilles Institute of Biodiversity and Evolutionary Biology, University of Valencia, Burjassot, E-46100 Valencia, Spain

**Keywords:** photic zone, deep chlorophyll maximum, Mediterranean, stratification, stenobathic

## Abstract

Most marine metagenomic studies of the marine photic zone analyze only samples taken at one or two depths. However, when the water column is stratified, physicochemical parameters change dramatically over relatively short depth intervals. We sampled the photic water column every 15m depth at a single point of an off-shore Mediterranean site during a period of strong stratification (early autumn) to evaluate the effects of small depth increases on the microbiome. Using genomic assembly and metagenomic read recruitment, we found major shifts in the community structure over small variations of depth, with most microbes showing a distribution limited to layers approximately 30 meters thick (stenobathic). Only some representatives of the SAR11 clade and the Sphingomonadaceae appeared to be eurybathic, spanning a greater range of depths. These results were confirmed by studying a single gene (rhodopsin) for which we also found narrow depth distributions. Our results highlight the importance of considering vertical distribution as a major element when analyzing the presence of marine clades and species or comparing the microbiome present at different locations.

## INTRODUCTION

The open ocean is one of the largest and most biologically productive microbial habitats in the biosphere. However, it is far from a homogenous habitat and environmental conditions are strongly affected by depth in the water column (Delong *et al.*, 2006; Konstantinidis *et al.*, 2009). As depth increases, many environmental factors change. Among them, light attenuation is of paramount importance since the availability of light is critical for primary productivity and hence is the main limiting factor for organic matter production throughout the water column (Letelier *et al.*, 2004). However, many other parameters change with depth, particularly temperature, salinity and inorganic nutrient availability (mostly nitrate and phosphate, although others such as iron or silicate can play a significant role for some biological groups). The main divide in aquatic environments tends to be between the photic zone, where light allows photosynthesis, and the aphotic, beyond the compensation depth, where the available light (if any) is insufficient to drive photosynthesis. The differences of the microbiome between the photic and aphotic zone have been extensively illustrated and analyzed by multiple approaches (Delong *et al.*, 2006; Ferreira *et al.*, 2014; Walsh *et al.*, 2015; Shi *et al.*, 2011). In the same way, global ocean scale studies such as those derived from the Sorcerer II Global Ocean Sampling (GOS) (Rusch *et al.*, 2007) or the recent Tara Oceans expedition (Sunagawa *et al.*, 2015) have provided essential baseline information to explore the spatial profile of the surface ocean microbial communities on a global scale but not the vertical distribution throughout the water column. In these studies, the photic zone, which is the most biologically relevant, is mostly considered to be a homogeneous single ecological compartment and what is found out for a single depth often considered representative of the complete photic water column. However, most of the off-shore oceanic waters are permanently or seasonally stratified creating strong shifts of environmental parameters that affect epipelagic communities inhabiting the photic zone proper (Cram *et al.*, 2014; López-Pérez *et al.*, 2016).

The Mediterranean is a nearly closed sea that represents a model of the global ocean. The water column is seasonally stratified, typically from March to November. A characteristic and extensively studied phenomenon associated with this stratification is the formation of the deep chlorophyll maximum (DCM) (Christaki *et al.*, 2011), a maximum in chlorophyll concentration that associated with an increase in bioavailable pools of N and P diffusing from the mixed layer below the seasonal thermocline (Denaro *et al.*, 2013). In the Mediterranean, the DCM appears between 45 and 70 m deep (Estrada *et al.*, 2004) depending on the light penetration (season of the year and water biological productivity). The water column is fully mixed during the winter due to the relatively warm deep water mass (*ca*. 13°C) and the absence of a permanent thermocline (Estrada *et al.*, 2004; Ghai *et al.*, 2010). The Mediterranean is oligotrophic overall because it has a negative hydric balance (more evaporation than fresh water inputs) and the incoming flow of Atlantic surface water through the Gibraltar straits rim is compensated by the deep outward current that pumps nutrients back into the deep Atlantic (Harmelin and Dhondt, 1993).

The majority of studies are focused on surface samples or DCM alone (Delong *et al.*, 2006; Rusch *et al.*, 2007; Ghai *et al.*, 2010) and a detailed profile of the stratified water column using the power of high throughput metagenomics is missing to date. Furthermore, in some previous studies, it is not guaranteed that the water column was stratified at the time of sampling (Sunagawa *et al.*, 2015). There are still many knowledge gaps regarding the microbes, particularly the prokaryotic ones that occupy the photic zone. This study is the first to use high throughput metagenomics to analyse a fine-scale profile (every 15m) of microbial communities in a stratified and mature (early autumn) Western Mediterranean water column. We detected marked stratification at the level of large clades, but mostly at finer taxonomic levels (species), that reflect adaptation of microbial species to live at defined ranges of depth in the water column.

## METHODS

**Sampling, sequencing, assembly and annotation**. Seven samples from different depths were taken for metagenomic analyses on 15^th^ October 2015 at a single site from the Western Mediterranean (37.35361°N, 0.286194°W), approximately 60 nautical miles off the coast of Alicante, Spain from the research vessel “García del Cid”. Six seawater samples (200 L each) were collected from the uppermost 100 m at 15 m intervals using a hose attached to a CTD (Seabird) deployed without the rosette and connected to a water pump in order to directly transfer seawater from selected depth to the filtration system and thus minimize sample storage time and potential bottle effects (Supplementary Figure 1). Each sample was filtered in less than 30 minutes. Another sample from a depth of 1000 m, was taken the next day (16^th^ October) in two casts (100 L each) using the CTD rosette. For comparison, another sample from a depth of 60 m was collected on September 12, 2015 from a location closer to the coast, approximately 20 nautical miles off the coast of Alicante (38.06851°N, 0.231994°W; bottom depth of 200 m), using the direct filtration method described above. Purposefully, samples were not collected from the surface since they are known to be inherently patchy spatially and temporally due to the hydrodynamic action of wind and currents, and can be affected by the presence of the ship used for sampling.

All seawater samples were sequentially filtered on board through 20, 5 and 0.22 μm pore size polycarbonate filters (Millipore). All filters were immediately frozen on dry ice and stored at −80°C until processing. DNA extraction was performed from the 0.22 μm filter as previously described (Martin-Cuadrado *et al.*, 2008). Metagenomes were sequencing using Illumina Hiseq-4000 (150 bp, paired-end read) (Macrogen, Republic of Korea). Individual metagenomes were assembled using IDBA-UD (Peng *et al.*, 2012). The resulting genes on the assembled contigs were predicted using Prodigal (Hyatt *et al.*, 2010). tRNA and rRNA genes were predicted using tRNAscan-SE (Lowe and Eddy, 1996), ssu-align (Nawrocki, 2009) and meta-rna (Huang *et al.*, 2009). Predicted protein sequences were compared against NCBI NR, COG (Tatusov *et al.*, 2001) and TIGFRAM (Haft *et al.*, 2001) databases using USEARCH6 (Edgar, 2010) for taxonomic and functional annotation. GC content, average genome size and richness in each sample were calculated using the gecee program from the EMBOSS package (Rice *et al.*, 2000), MicrobeCensus (Nayfach and Pollard, 2015) and MEGAN6 Community Edition (Huson *et al.*, 2016), respectively.

**Environmental metadata**. Vertical profiles of temperature and chlorophyll-*a* concentration were determined *in situ* using a temperature profiler including a fluorometer (Seabird). Analyses of chemical variables and microbial counts were performed on the same samples retrieved for the metagenomics analyses. Chemical analyses included the measurement of inorganic soluble forms of nitrogen (NOx and ammonium) and phosphorus (soluble reactive phosphorus), and of total nitrogen (TN) and total phosphorus (TP), all of them following standard analytical methods (Eaton *et al.*, 1992). Total organic carbon (TOC) was also determined on acidified samples using a Shimadzu TOC-VCSN Analyzer. To refine the results given by the fluorometric probe used in situ, photosynthetic pigments were further determined by HPLC on a Waters HPLC system after filtration of the samples onto GF/F filters and extraction in acetone (Picazo *et al.*, 2013).

**Microbial counts**. A Coulter Cytomics FC500 flow cytometer (Brea, California, USA) equipped with an argon laser (488 excitation) and a red emitting diode (635 excitation) was used to perform flow cytometric counts of picoplankton samples. Analyses were run for 160 seconds at the highest possible single flow rate (128μL min^−1^). Data were collected using the Beckman Coulter software for acquisition “CXP Version 2.2 Acquisition” and the analysis of the data was performed using Beckman Coulter software for analysis “CXP Version 2.2 Analysis”. Relative DNA content was determined by Sybr Green I (Sigma-Aldrich, Missouri, USA) fluorescence at 525 nm. The abundance of autotrophic picoplankton was determined by measuring chlorophyll a and phycobiliprotein autofluorescence excited by the argon laser and the red diode. *Synechococcus* and *Prochlorococcus* cells were differentiated by both their fluorometric and size features as previously described (Marie *et al.*, 1997).

**Phylogenetic classification**. A non-redundant version of the RDP database (Cole *et al.*, 2014) was prepared by clustering all available 16S/18S rRNA gene sequences (*ca*. 2.3 million) into approximately 800,000 clusters at 90% identity level using UCLUST (Robert C Edgar, 2010). This database was used to identify candidate 16S/18S rRNA gene sequences in the raw metagenomes (subsets of 10 million reads). Using USEARCH, sequences that matched this database (e-value < 10^−5^) were considered potential 16S rRNA sequences. These candidates were then aligned to archaeal, bacterial and eukaryal 16S/18S rRNA HMM models (Eddy, 1995) using ssu-align to identify true sequences (Nawrocki, 2009). Final 16S/18S rRNA sequences were compared to the entire RDP database and classified into a high level taxon if the sequence identity was ≥80% and the alignment length ≥90 bp. Sequences failing these thresholds were discarded.

**Binning and genome reconstruction**. Assembled contigs longer than 10 kb were assigned a high level taxon classification if >50% of the genes shared the same taxonomy (Supplementary Figure 3). Contigs were then binned together using principal component analysis of tetranucleotide frequencies, GC content and coverage values within their source metagenome. Tetranucleotide frequencies were computed using wordfreq program in the EMBOSS package (Rice *et al.*, 2000). Principal component analysis was performed using the FACTOMINER package (Lê *et al.*, 2008) in R. Completeness of the genomes was estimated by comparison against two different universal gene sets, one with 35 genes (Raes *et al.*, 2007) and another with 111 genes (Albertsen *et al.*, 2013). Additionally, the degree of contamination was estimated with CheckM (Parks *et al.*, 2015). In order to improve the completeness and remove the redundancy, a second assembly step was performed combining the genomic fragments with the short paired-end Illumina reads of the metagenomes from they were assembled. For each genome, we used the BWA aligner (Li and Durbin, 2009) with default parameters to retrieve the short paired reads that mapped onto the contigs. These reads were then pooled and assembled together with the contigs using SPAdes (Bankevich *et al.*, 2012).

**Metagenomic read recruitments**. The genomes of known marine microbes along with genomes reconstructed in this study were used to recruit reads from our metagenomic datasets using BLASTN (Altschul *et al.*, 1997), using a cut-off of 99% nucleotide identity over a minimum alignment length of 50 nucleotides. Genomes that recruited less than three reads per kilobase of genome per gigabase of metagenome (RPKG) were discarded.

**Phylogenomic trees of the reconstructed genomes**. Phylogenomic analysis was used to classify and identify the closest relatives for all reconstructed genomes. Using HMMscan, we aligned the sequences against the COG database. Shared proteins were concatenated and aligned using Kalign (Lassmann and Sonnhammer, 2005). A maximum-likelihood tree was then constructed using MEGA 7.0 (Kumar *et al.*, 2016) with the following parameters: Jones-Taylor-Thornton model, gamma distribution with five discrete categories, 100 bootstraps. Positions with less than 80% site coverage were eliminated.

**Metagenomic cross-comparisons**. Two different approaches were used to compare similarities between metagenomic samples. First, a reciprocal global alignment of the short Illumina reads (in subsets of 2 million of reads ≥ 50 bp) at ≥ 95% identity was performed using USEARCH6 (Edgar, 2010). Results of the comparison were then clustered with the hclust package in R using an euclidean distance matrix. In a second approach, subsets of 20 million of reads ≥ 50 bp (where applicable) were taxonomically classified against the NR database using DIAMOND (Buchfink *et al.*, 2015) with a minimum of 50% identity and 50% alignment. The resulting alignment was later analyzed with MEGAN6 Community Edition (Huson *et al.*, 2016), and a canonical correspondence analysis (CCA) was inferred with the cluster analysis option and a Bray-Curtis ecological distance matrix.

**Rhodopsin.** 168 rhodopsin sequences were extracted from all the metagenomes from assembled contigs longer than 5Kb. These sequences were pooled with one hundred more rhodopsins of fungal, archaeal, viral and bacterial origin obtained from databases. Sequences were aligned with MUSCLE (Edgar, 2004) and a Maximum-Likelihood tree was constructed with MEGA 7.0 (Jones-Taylor-Thornton model, gamma distribution with five discrete categories, 100 bootstraps, positions with less than 80% site coverage were eliminated). Blue versus green light absorption was determined as described previously (Man *et al.*, 2003). To compare the abundance of microbial rhodospins with depth, we initially created a database containing our metagenomic rhodopsin sequences and approximately 7900 rhodopsin genes obtained from the MicRhoDE database (http://micrhode.sb-roscoff.fr). Metagenomic reads (in subsets of 20 million sequences) were recruited to these rhodopsin sequences using BLASTN (≥50bp alignment, ≥99% identity). Rhodopsin sequences that recruited ≥ 1RPKG were kept for further analyses. In parallel, metagenomic reads were compared to the NR database using DIAMOND (blastx option, top hit, ≥ 50% identity, ≥ 50% alignment length, e-value < 10^−5^). The abundance of rhodopsin genes in each metagenome was estimated from the number reads matching rhodopsin sequences in NR, normalized by the number of reads matching *recA/radA* sequences. Reads matching viral or eukaryal proteins were not taken into account.

**SAR11 clade analyses.** All the available *Ca*. Pelagibacter genomes were downloaded from the NCBI database, but only those longer than 300 Kb were selected. Average nucleotide identities (ANI) and average amino acid identity (AAI) among SAR11 representatives were calculated using BLASTN and CompareM (https://github.com/dparks1134/CompareM) respectively, with thresholds of 50% (60% for ANI) identity and 50% alignment. SAR11 genomes were grouped into clusters based on the ANI and AAI results. The abundance of the SAR11 genomes was estimated using established approaches (Tsementzi *et al.*, 2016). Briefly, a reference protein database was built containing all the available SAR11 genomes (52). To this database, we added predicted proteins from all the reference genomes used for metagenomic recruitments in this study (385), as well as all predicted proteins from our assembled contigs ≥10 Kb that were not taxonomically classified as Alphaproteobacteria (6,871 contigs). Potential SAR11 reads were extracted from the metagenomic reads using DIAMOND (blastx option, top hit, ≥50% identity, ≥50% alignment length, evalue < 10^−5^), thus defining the fragments that can be considered bona fide Pelagibacterales. The number of orthologous genes (OG) among these clusters was identified using rbm.rb and ogs.mcl.rb from the enve-omics collection (Rodriguez-r and Konstantinidis, 2016). Then, the potential Pelagibacter reads were used to quantify the abundance of each clade within the metagenomes. Using DIAMOND (blastx option, top hit, ≥50% identity, ≥50% alignment length, evalue <10^−5^), best hits against the OGs of each subcluster were counted as a recruited item. The value was normalized by gene size (bp) and number of Pelagibacter reads for each metagenome.

**Data availability.** Metagenomic datasets have been submitted to NCBI SRA, and are available under BioProjects accession number PRJNA352798 (Med-OCT2015-15m, Med-OCT2015-30m, Med-OCT2015- 45m, Med-OCT2015-60m, Med-OCT2015-75m, Med-OCT2015-90m, Med-OCT2015-1000m and Med-OCT2015-2000m) and PRJNA257723 (Med-SEP2015_HS). The reconstructed genomes have been deposited as BioSample NNNNNNNXXX to NNNNNNXXX under BioProject PRJNA352798.

## RESULTS

Seawater samples from six depths in the photic zone were collected at 15 m intervals (15, 30, 45, 60, 75 and 90 m) on a single day. For comparison, we also collected another two samples: one from 1000 m from the same site as above and another sample from 60 m at a different site. With the exception of the 1000 m sample, all samples were collected using a hose directly connected to the filtration apparatus to minimize processing time and to avoid bottle effects (Supplementary Figure 1). Metadata and sequencing results are described in Table 1.

**Table 1.** Summary statistics of the sampling, sequencing and assembly parameters.

|  | Med-OCT2015-15m | Med-OCT2015-30m | Med-OCT2015-45m | MedDCM-OCT2015-60m | Med-OCT2015-75m | Med-OCT2015-90m | Med-OCT2015-1000m | MedDCM-SEP2015 |
| --- | --- | --- | --- | --- | --- | --- | --- | --- |
| **Sampling parameters** |  |  |  |  |  |  |  |  |
| Sampling data |  |  |  | 10/15/2015 |  |  |  | 9/12/2015 |
| Depth (m) | 15 | 30 | 45 | 60 | 75 | 90 | 1000 | 60 |
| Sea bottom depth (m) |  |  |  | 2647 |  |  |  | 200 |
| Size fraction |  |  |  | 5-0.22 µm |  |  |  | 5-0.22µm |
| Latitude |  |  |  | 37.35361 N |  |  |  | 38.06851 N |
| Longitude |  |  |  | 0.286194 W |  |  |  | 0.231994 W |
| **Environmental parameters** |  |  |  |  |  |  |  |  |
| Temperature (°C) | 22.90 | 18.40 | 15.80 | 14.50 | 14.00 | 13.80 | 13.10 | 14.60 |
| Chl-a (mg/m<sup>3</sup>) | 0.14 | 0.21 | 0.37 | 0.54 | 0.26 | 0.13 | 0.05 | 0.80 |
| Oxygen (mg/L) | 7.09 | 8.77 | 9.00 | 7.66 | 7.16 | 6.88 | 6.10 | 6.62 |
| TOC (mgC/L) | 2.43 | 2.17 | 1.46 | 1.43 | 1.36 | 1.35 | 0.84 | 1.12 |
| PO<sub>4</sub><sup>3-</sup> (µM) | 0.06 | 0.07 | 0.10 | 0.08 | 0.12 | 0.22 | 0.39 | 0.09 |
| Total P (µM) | 0.10 | 0.12 | 0.14 | 0.12 | 0.16 | 0.25 | 0.45 | 0.13 |
| NO<sub>x</sub> (µM) | 0.20 | 0.21 | 0.25 | 0.23 | 5.79 | 6.23 | 8.24 | 0.25 |
| NH<sub>4</sub><sup>+</sup> (µM) | 0.13 | 0.12 | 0.14 | 0.15 | 0.09 | 0.08 | 0.03 | 0.09 |
| Total N (µM) | 0.40 | 0.41 | 0.48 | 0.46 | 6.33 | 6.90 | 8.89 | 0.39 |
| N:P ratio | 4.00 | 3.42 | 3.43 | 3.83 | 39.56 | 27.60 | 19.76 | 3.00 |
| **Sequencing statistics** |  |  |  |  |  |  |  |  |
| Total sequenced (Gb) | 19.9 | 15.1 | 3.1 | 15.3 | 16.9 | 15.0 | 14.8 | 20.0 |
| Mean read length (bp) | 121.4 | 117.2 | 112.3 | 119.8 | 121.4 | 120.2 | 117.0 | 139.9 |
| Mean GC (%) | 38.6 | 41.4 | 40.4 | 40.6 | 41.3 | 41.1 | 45.9 | 38.7 |
| **Assembly statistics** |  |  |  |  |  |  |  |  |
| Total assembled (Mb) | 738.5 | 500.3 | 79.5 | 577.3 | 701.3 | 613.5 | 490.0 | 916.1 |
| Mean GC (%) | 38.5 | 35.7 | 35.5 | 38.6 | 38.9 | 40 | 42.2 | 35.3 |
| Maximum contig length (Kb) | 251 | 165 | 118 | 235 | 186 | 140 | 450 | 494 |
| Contigs > 1Kb | 172,484 | 125,642 | 17,760 | 145,031 | 173,153 | 153,245 | 115,711 | 230,853 |
| Contigs > 10Kb | 4,648 | 2,198 | 202 | 2,360 | 2,789 | 2,556 | 1,807 | 4,538 |

### Variability of environmental parameters

Because of the prolonged physical isolation and accumulation of settled organic matter during the summer stratification period, bottom waters are richer in both dissolved and particulate inorganic nutrients (N, P) (Table 1). However, layers within the DCM are typically the richest of the photic zone both in term of biomass accumulation and N and P availability. Chlorophyll-*a* measurements indicated that the DCM occurred between 40-60 m depth, just below the seasonal thermocline (Figure 1a). Chlorophyll-*a* measurements reached 0.8 mg•m^-3^, almost one order of magnitude above those from surface waters and hundred times those of deep (1000m) waters. Absolute maximal abundances of planktonic picoprokaryotes for the whole water column, especially *Prochlorococcus* (nearly 3.2 ×10^4^ cells mL^-1^) and, to a lesser extent, also *Synechococcus* (1.35 ×10^4^ cells mL^-1^), were found in the DCM peak. These values were 1-2 orders of magnitudes higher than in surface waters. The distribution of *Phrochlorococcus* cells was wider than that of *Synechococcus* (Figure 1a), as previously described (Partensky *et al.*, 1999).

**Figure 1.**
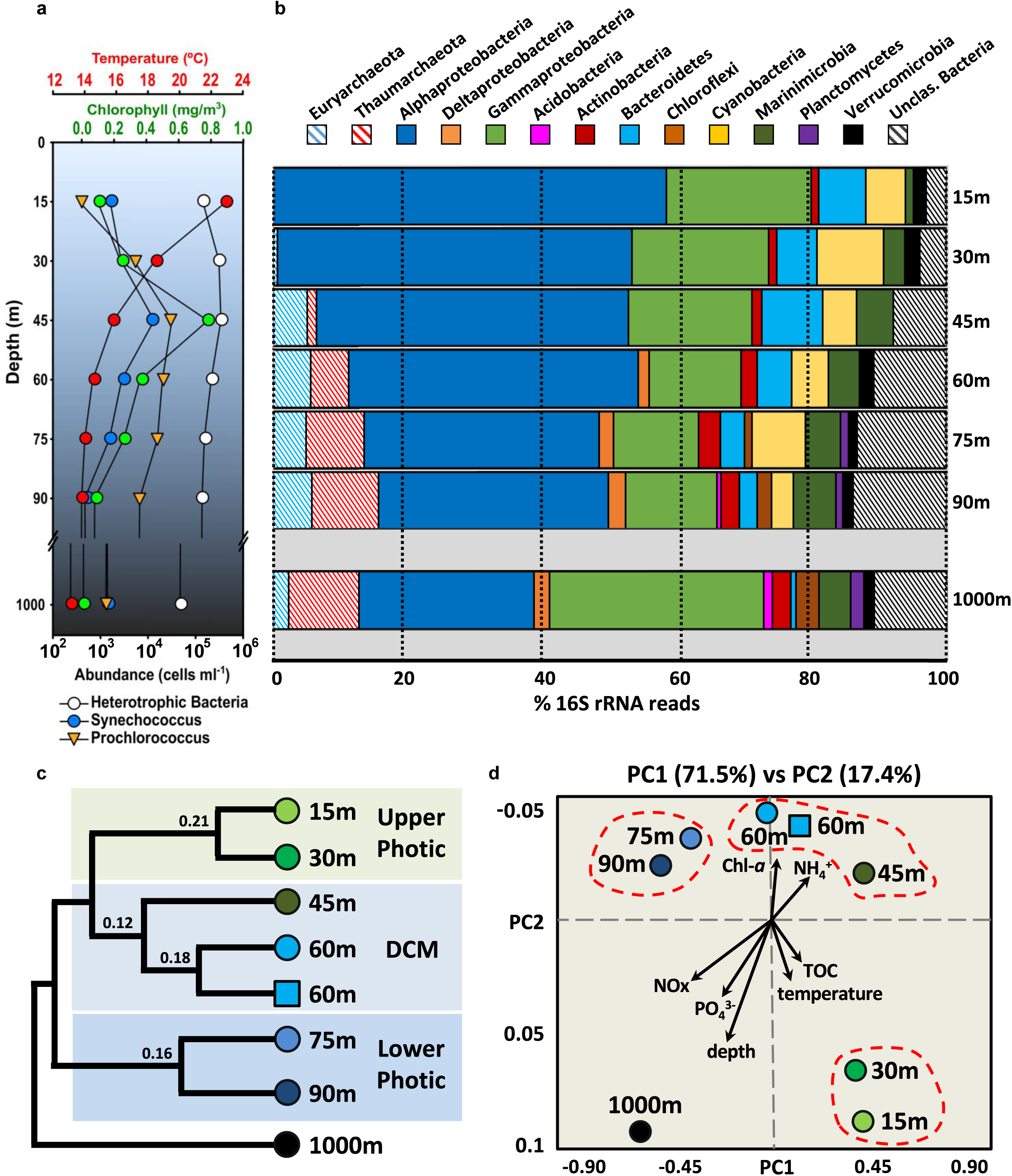
a) Vertical profile of the temperature and chlorophyll *a* measurements, along with the abundance of heterotrophic bacteria, *Synechococcus* and *Prochlorococcus* representatives. b) Class level composition based on 16S rRNA (raw reads) of the different depth metagenomes. c) Dendrogram of metagenomic datasets raw sequence similarity. Sample represented by a blue square belongs to a different sampling site closer to the coast. d) Canonical correlation analysis between physicochemical parameters and read annotation similarity. Chl-a: chlorophyll-a, TOC: total organic carbon.

### Overall community structure and depth

Using the percentage identity of individual reads between metagenomes, we examined the relationship between the eight sequenced samples (Figure 1c). As expected, samples clustered by depth, with three main branches corresponding to i) upper photic (UP, 15 and 30 m), ii) DCM (45 and 60 m) and iii) lower photic (LP, 75 and 90 m). A separate sample from 60m depth collected from the DCM in a comparable environment closer to the coast (20 instead than 60 nautical miles, bottom depth 200 m) one month before the vertical profile clustered with the off-shore 60m depth sample. As expected the bathypelagic 1000 m sample appeared as an outgroup to all photic zone samples. Independently, canonical correlation analysis (CCA) of read annotations and environmental parameters confirmed the clustering of samples according to depth (Figure 1d). Inorganic nutrients (e.g., NOx and PO_4_^3-^) increased with depth, while ammonia correlated closely with chlorophyll *a*, and total organic carbon (TOC) increased at the surface together with water temperature. These results clearly indicate that depth is a greater driver of community composition than distance from shore.

General genomic parameters also varied with depth (Supplementary Figure 2). GC content and average genomes size were smaller at 15 m (*ca*. 38.2% and 2.5 Mb) and higher at 1000 (*ca*. 44.5% and 3.8 Mb), while remaining relative constant throughout the photic zone deeper than 15m (ca. 42% and 3.5 Mb). The lower GC content observed in near-surface stratified waters has been suggested to be a natural adaptation to reduce N demand in these environments with severe depletion of bioavailable N pools (Luo *et al.*, 2015). In the photic zone, average genome size was greatest in the DCM (approximately 4 Mb at 45 m), decreasing to approximately to 3 Mb in the 90 m sample. However in the bathypelagic sample (1000 m), the average estimated genome size increased again to approximately 4 Mb. Measurements of Simpson’s Diversity Index at both genus and species level clearly indicated that bacterial diversity increased continuously with depth (Supplementary Figure 2). The 15 m sample was the least diverse of the photic zone, with a marked domination of Pelagibacterales. At this depth, high light intensity and nutrient depletion generate conditions that can be considered extreme and that may result in the survival of only a few microbial taxa. In deeper waters, diversity increased with depth, particularly at the species level, reaching a maximum at our deepest sample at 1000 m (Supplementary Figure 2). The larger diversity and genome size of microbes in bathypelagic waters might correlate with a capacity to degrade or use a larger number of different substrate compounds.

Metagenome derived 16S rRNA profiles revealed broad, depth-dependent variations in taxonomic range (Figure 1b). Archaea, absent in the UP region, represent nearly 16% of the population at 90 m. In the DCM and LP samples, Euryarchaeota remained constant (*ca*. 5%), while Thaumarchaeota increased from 1% in the 45 m sample to 10% of all the rRNA reads in the 90 m sample. The abundance of Thaumarchaeota correlated with a sharp decrease in ammonia concentration, although the main increase in Thaumarchaeota occurred at 60 m while ammonia concentrations decreased most at 75 m (Figure 1 and Table 1). Whereas Actinobacteria, Bacteroidetes, Cyanobacteria and Marinimicrobia were present in the whole water column, Deltaproteobacteria, Planctomycetes, Chloroflexi and Acidobacteria had a much more restricted range, appearing in deeper layers of the photic zone (Figure 1). Interestingly, Verrucomicrobia were present at all depths except the 45 m sample (Figure 1b). Using finer-scale taxonomic classification of the 16S rRNA sequences, we found that UP (15 and 30 m) Verrucomicrobia belonged to the *Puniceicoccaceae*, whereas members of the *Verrucomicrobiaceae* were found predominantly below the DCM (Supplementary Table 1). The proportion of 16S rRNA reads assigned to unclassified bacteria also increased with depth, from 3% at 15 m to more than 10% at 90m, indicating that a significant fraction of the microbes at the subsurface are still uncharacterized.

### Fine taxonomic profile: genome recruitments

Unfortunately, the broad organismal distributions detected by 16S rRNA genes or raw sequence annotation methods described above do not shed light on more subtle, but ecologically significant, variations of community structure or metabolic function that likely occur at the finer levels of diversity, such as species, ecotypes or even clonal lineages within species (Biller *et al.*, 2014; Kashtan *et al.*, 2014; Bendall *et al.*, 2016; Gonzaga *et al.*, 2012). To investigate the distribution of major ecotypes present in the water column, we used stringent recruitment of metagenomic reads to assemblies of locally predominant genomes. In general, genome assembly improved proportionally to the abundance of the phylum. However, we found that genomes of representatives from the Bacteroidetes, Actinobacteria and Acidobacteria assembled better than expected (Supplementary Figure 3). On the other hand, Cyanobacteria and Thaumarchaeota assembled much more poorly. These differences in ease of genome assembly are likely driven by variations in the amount of intraspecies diversity and intragenomic heterogeneity. As such, we obtained very few long contigs belonging to either the Pelagibacterales or the picocyanobacteria, despite the abundance of reads assigned to both groups (Figure 1b). Both groups are known to possess enormous intra-species diversity (Grote *et al.*, 2012), explaining why assembly for these two major components of the bacterioplankton was very poor. Genome assembly improves when individually assembled clones have very different genomes or when small numbers of clones predominate. In contrast, intragenomic redundancy (e.g., multiple copies of rRNA genes or IS elements) would complicate assembly. However, redundancy is expected to be low in oligotrophic marine environment that tend to be dominated by streamlined genomes.

Overall, we have retrieved 44 genomes belonging to those phyla in which we obtained more than 8Mb of assembled contigs (Supplementary Table 2 and Supplementary Figure 3), using a combination of different parameters such as GC content, metagenomics read coverage, and tetranucleotide frequencies. These genomes were classified phylogenomically using concatenated sequences of conserved proteins (Figure 2 and Supplementary Figures 4-10). Additionally, Figure 2 shows the recruitment of the assembled genomes plus several reference genomes from public databases that recruited at least three RPKG of coverage with a similarity >99% in our metagenomic profile. The complete list of the 74 genomes and their recruitment values is given in Supplementary Table 3.

**Figure 2.**
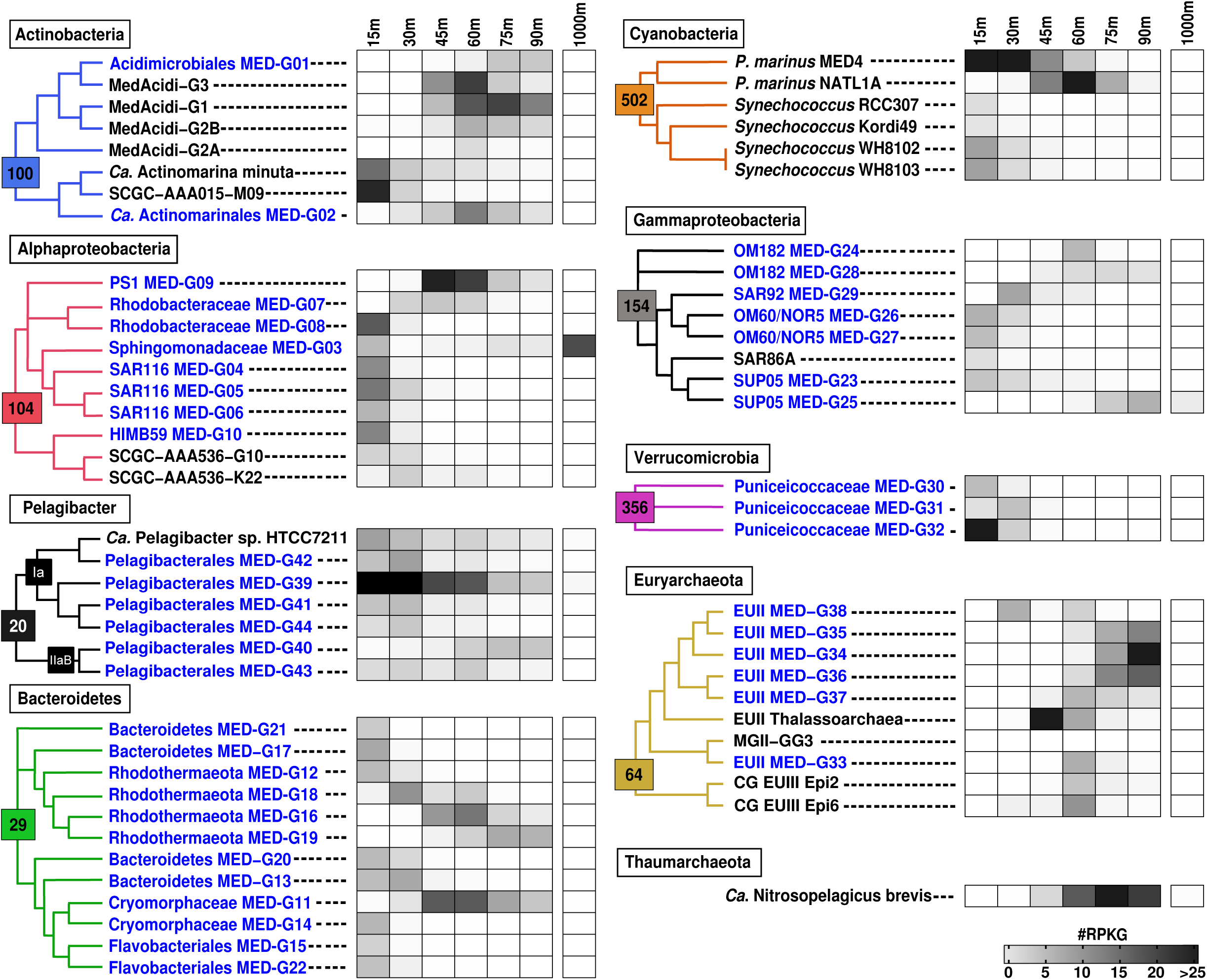
a) Relative abundance of the reconstructed and reference genomes measured by recruitment (RPKG, Reads per Kilobase of genome and Gigabase of metagenome) from the different depths metagenomes. To show the relationships among genomes, a maximum likelihood genome tree was constructed using all the conserved proteins (number in coloured square). Each reconstructed genome (in blue). has been assigned a name derived from their position in the phylogenomic tree built with the closest known relatives from databases and presented in the Supplementary figures 4 to 9). Black genomes are from previously reported works (cultures, SAGs or reconstructed genomes from metagenomes)

### Actinobacteria

We reconstructed two genomes belonging to the bacterial phylum Actinobacteria from our metagenomes, MED-G01 belonging to the subclass Acidimicrobidae (Mizuno *et al.*, 2015) and MED-G02 belonging to the subclass “*Ca.* Actinomarinidae” (Ghai *et al.*, 2013) (Supplementary Figure 4). MED-G01 (1.96Mb, 53.1% GC content) was most closely related to MedAcidi-G3 (2.1Mb, 50-52%GC content), a genome previously reconstructed from a Mediterranean metagenome (Mizuno *et al.*, 2015). These two genomes, alone with four additional reconstructed genomes from the same study (Mizuno *et al.*, 2015), recruited significant numbers of reads (RPKG ≥ 3) from our photic zone datasets, predominantly in the DCM (45, 60 and 75 m). MedAcidi-G1 showed a wider distribution down to 90 m; whereas MedAcidi-G2A was found exclusively at 60 m. MED-G02 belongs to the subclass “*Ca*. Actinomarinidae” (72.9% ANI) that is widely distributed in the photic zone in marine waters (Ghai *et al.*, 2013). However, while *Ca*. Actinomarina minuta, the original representative reconstructed from a DCM fosmid library, and the single amplified genome (SAG) SCGC-AAA015-M09 recruited the most at 15 m decreasing with depth (Figure 2), MED- G02 showed a clear maximum at 60 m.

### Alphaproteobacteria

Alphaproteobacteria represented the largest lineage in our metagenomic dataset, particularly in the photic zone (Figure 1). The more than 50Mb of assembled contigs affiliated with Alphaproteobacteria fell into eight different bins and, based on a phylogenomic tree of concatenated proteins, were classified into five distinct clades: PS1, Rhodobacterales, Sphingomonadales, SAR11 and SAR116 (Supplementary Figure 5). MED-G09 clustered with the reference genomes of the PS1 clade (Jimenez-Infante *et al.*, 2014), RS24 and IMCC14465. Although these strains were cultivated from a surface seawater sample, recruitment analysis showed that MED-G09 was found only at the DCM and at much lower levels deeper in the photic zone (Figure 2). MED-G07 and G08 were associated with members of the family *Rhodobacteraceae* within the ubiquitous and versatile Roseobacter clade (Supplementary Figure 5). These genomes contained genes for bacteriochlorophyll-based aerobic anoxygenic phototrophy, a common feature of this group. However both genomes showed different depth preferences, with MED- G07 found exclusively at 15 m, and MED-G08 most abundant for 30 to 60 m. MED-G03 belongs to the family *Sphingomonadaceae* that have been characterized by the ability to degrade a broad range of polycyclic aromatic hydrocarbons and polysaccharides (Balkwill *et al.*, 2006). This was the largest genome assembled with 3.15Mb. MED-G03 was most closely related to *Sphingomonas* sp. SKA58 and *Sphingobium lactosutens*, and had an ANI of *ca*. 84% to both of these genomes, consistent with a new species within the genus. Although MED-G03 consisted of contigs from the 15 m metagenome, recruitment analysis indicated that it was most abundant at 1000 m (RPKG= 19). MED-G04, G05 and G06 belonged to the SAR116 clade, a ubiquitous group of heterotrophic bacteria inhabiting the surface of the ocean (Giovannoni and Vergin, 2012). All three genomes were restricted to the 15 m sample and were most closely related to the HIMB100 and “*Candidatus* Puniceispirillum marinum” IMCC1322 genomes (Supplementary Figure 5). MED-G10 clustered together with HIMB59 (Supplementary Figure 5), a pure culture often considered representative of the SAR11 subclade V but which is only distantly to other members of the SAR11 clade (Viklund *et al.*, 2013).

### SAR11 subclade of Alphaproteobacteria

Based on rRNA data, the most abundant clade was SAR11. However, as mentioned above, assembly of SAR11 genomes from our metagenomes was very poor. Furthermore, the SAR11 genomes available in public databases were very different from the contigs assembled here. We were able to assemble six genomes but using only contigs coming from LP metagenomes were the genomic variability was likely smaller (Supplementary Table 2). To estimate the abundance of different SAR11 subclades we used 46 available SAR11 genomes and our six assemblies to conduct recruitment analysis using previously described methodology (Tsementzi *et al.*, 2016). We performed a new reclassification of the SAR11 subclades based on pairwise comparisons using both average amino acid identity (AAI) and average nucleotide identity (ANI) distances, obtaining 11 different subclades (Supplementary Figure 11). Subtype Ic is thought to be most prevalent in deep waters (Thrash *et al.*, 2014) and recruited more than 40% of the SAR11 reads at 1000m (Figure 3). Although, the remaining SAR11 subtypes were widespread throughout the entire water column and could be considered eurybathic (Figure 3), many appeared to be most abundant at discreet depths. For instance subclade IaA was found predominantly in the DCM and LP depths (below 45 m), subclade IaD was most abundant at the DCM and subclade IaC appeared to prefer UP (15 and 30 m). Members of the subclades IaB and IaE together with IIaB were the most prevalent in all depths while subclades IIaA, III and V were the least abundant. The reconstructed genome Pelagibacterales MED-G39, belonging to subclade IaC (Supplementary Figure 11), showed the highest recruitment value of all the microbes analyzed throughout in the water column (RPKG=398 at 15 m). Furthermore, this genome recruited the most from all Mediterranean surface metagenomes but not at all from surface metagenomes from other ocean provinces (TARA and our own; Supplementary Table 4). MED-G39 may represent the most abundant and endemic Mediterranean Pelagibacter since it was not present anywhere else.

**Figure 3.**
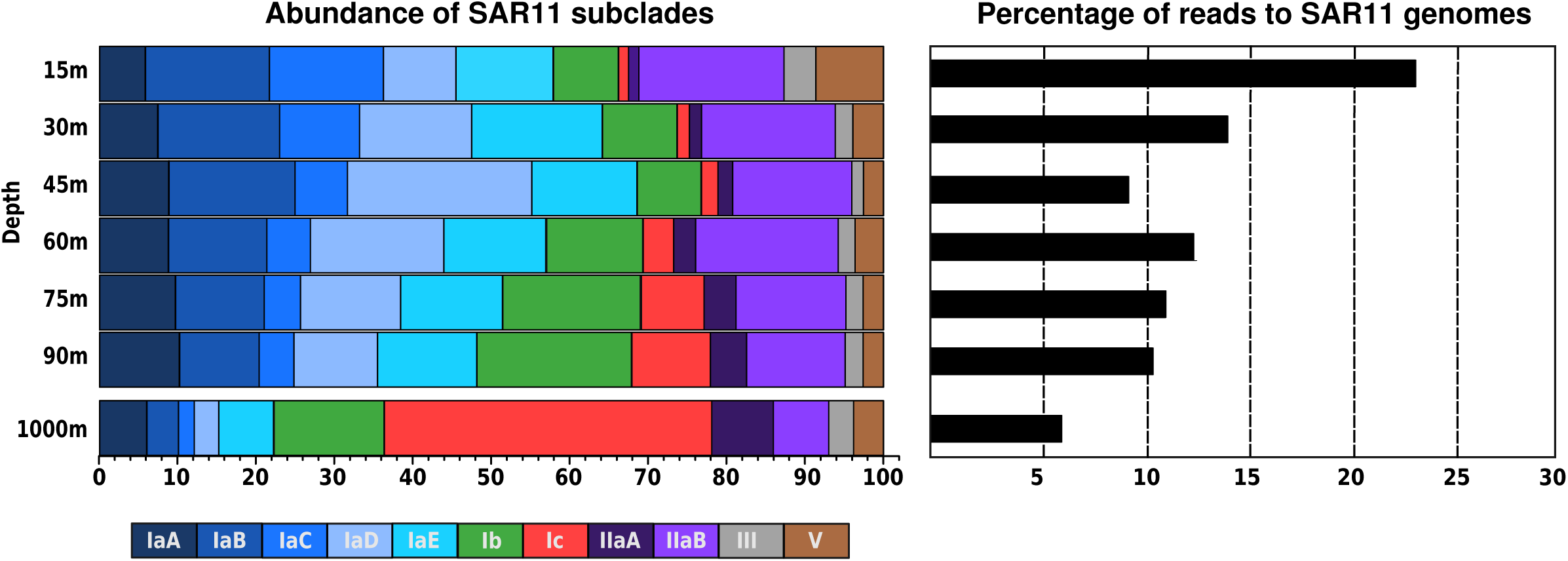
Relative abundance at different depths of SAR11 subclades. Percentage of reads belonging to the SAR11 clades in shown on the right panel.

### Bacteroidetes

Bacteroidetes was the third most abundant group in number of assembled contigs (1,209) and assemblies clustered into twelve new genomic bins, the highest number of any taxon in this study. Members of this group have been characterized as particle-attached microbes that degrade polymeric organic matter, mainly polysaccharides and proteins (Fernández-Gómez *et al.*, 2013). Phylogenomic analysis placed MED-G12, MED-G16, MED-G18 and MED-G19 genomes into the new proposed phylum Rhodothermaeota (Munoz *et al.*, 2016) (Supplementary Figure 6). MED-G12, with a GC content of 41%, clustered together with some members of the family Balneolaceae and was found only in the first 30 meters. MED-G16, MED-G18 and MED-G19 formed an independent new group with lower GC (*ca*. 30%), each with a distinct distribution pattern. MED-G18 was mainly localized at 30 m, whereas MED-G16 was found from 45 to 75m and MED-G15 between 75m and 90m. Four genomes, MED-G11, MED-G14, MED- G15 and MED-G22, were classified into the order Flavobacteriales. MED-G11 and MED-G14 belong to the family Cryomorphaceae, whereas MED-G15 and MED-G22 could represent a new family (Supplementary Figure 6). MED-G11 was widely distributed from 30 to 90 m. In contrast, the other Flavobacteriales genomes were found exclusively at 15m. The rest of the genomes retrieved designated only as “Bacteroidetes” belonged to new taxa (Supplementary Figure 6) since the average nucleotide and amino acid identities to the closest relatives was very low. They only recruited at the UP (15 and 30m). Genomic analysis of the reconstructed Bacteroidetes genomes indicated they encoded less metabolic diversity with an overrepresentation of polysaccharide degrading enzymes.

### Cyanobacteria

Cyanobacteria play an essential role in marine primary production (Field *et al.*, 1998) and carbon fixation (Zwirglmaier *et al.*, 2007) in the ocean. The vast majority of cyanobacterial sequences were classified into the genera *Prochlorococcus* and *Synechococcus* based on the annotation of our assembled contigs and rRNA analyses. The assembly of both groups was very poor (only 4.2 Mb in total). However, all contigs gave very high identity (>99%) to known genomes of cultivated representatives such as *Prochlorococcus* MED4 or NATL1A. We analyzed the presence of all 32 available genomes of cultivated representatives (23 *Synechococcus* and 9 *Prochlorococcus*) belonging to the different clades or ecotypes within each genus (Figure 2 and Supplementary Figure 7). Only six of genomes gave significant recruitment values in our metagenomes (Figure 2). Strains *Synechococcus* WH8102 and WH8103, motile members of the clade III, were dominant within this clade but were only detected at UP depths. Similar results were reported at a nearby sampling station (Balearic Sea, NW Mediterranean) using hybridization with clade-specific 16S rRNA oligonucleotide probes (Mella-Flores *et al.*, 2011). *Synechococcus* strains KORDI-49 and RCC307, belonging to subclusters 5.1 and 5.3-III respectively, exhibited a similar distribution pattern, although restricted only to the 15 m sample. In contrast, only two of the nine of available *Prochlorococcus* genomes recruited significantly from our metagenomes: MED4, a high-light (HLI)-adapted ecotype, and NATL1A, a low-light (LLI) ecotype. As expected based on prior studies (Garczarek *et al.*, 2007; Biller *et al.*, 2014), recruitments showed a clear stratification of MED4 and NATL1A along the profile. The first 45 m were dominated by *Prochlorococcus* MED4 (HLI), with a peak at 30 m, whereas *Prochlorococcus* NATL1A (LLI) predominated at DCM depths (Figure 2). In addition to their adaptation to light availability, the distribution of MED4 and NATL1A also correlated with their overall physiology (Tolonen *et al.*, 2006; Thompson *et al.*, 2011).

### Gammaproteobacteria

A total of 2,498 assembled contigs (the second largest number), clustering into seven distinct genomic bins, were assigned to the bacterial Gammaproteobacteria clade. Five of these genomic bins were affiliated with the oligotrophic marine gammaproteobacteria (OMG) group (Cho and Giovannoni, 2004) within the environmental clades SAR92, OM60/NOR5 and OM182. Phylogenomic analysis placed MED- G26 and G27 genomes in the OM60/NOR5 clade (Supplementary Figure 8), which contains aerobic anoxygenic photoheterotrophic bacteria commonly detected throughout the euphotic zone of marine environments especially in coastal waters (Yan *et al.*, 2009). These genomes showed low ANI values (*ca*. 70%) compared to the closest cultivated relative, HIMB55 (Supplementary Figure 8). MED-G26 contained a complete photosynthetic super-operon containing the cluster of genes related to bacteriochlorophyll biosynthetic pathway, subunits of the light-harvesting complex and carotenoid biosynthetic pathway. This gene cluster was only partially present in the close relative MED-G27. Aerobic anoxygenic photoheterotrophs play an important role in the ocean’s carbon cycle supplementing Chl*a*-based photosynthesis and contributing up to 5.7% of the total phototrophic energy, mainly in the surface layer, since they are not affected by photoinhibition (Jiao *et al.*, 2010). Accordingly, both genomes showed a similar distribution pattern restricted only to the UP (Figure 2a).

MED-G29, restricted to the 30 m sample, belonged to the ubiquitous gammaproteobacterial clade SAR92 represented by the cultivated strains HTCC2207 and MOLA455. Both strains contain a proteorhodopsin gene followed by an operon containing genes that are presumably involved in the synthesis of retinal. However, no proteorhodopsin gene could be found in MED-G29. MED-G24 and G28 were identified as a member of the OM182 group. However, while the presence of MED-G24 was restricted only to the DCM, MED-G28 showed a broader range of distribution (from 45 to 90m).

Members of the recently designated SUP05 clade of marine gammaproteobacterial sulfur oxidizers (GSOs) are among the most abundant chemoautotrophs in the ocean and are mainly found in low-oxygen environments (Glaubitz *et al.*, 2013; Walsh *et al.*, 2009). Phylogenomic analysis placed MED-G23 and G25 within the SUP05 clade (Supplementary Figure 8). Both genomes contain clusters of genes involved in chemotrophic sulfur oxidation including the dissimilatory sulfite reductase (*dsrAB*), the sulfur-oxidation cluster (*soxABXYZ*) or the adenylyl-sulfate reductase (*aprABM*). These are common features of the SUP05 group that provide the capability to reduce several sulfur compounds. G23 and G25 had low ANI (<70%) and displayed nearly opposite depth distributions, with MED-G23 present in the UP and MED- G25 in the LP. The presence of these organisms in oxygenated photic zone layers is at odds with prior reports of SUP05 distributions (Swan *et al.*, 2011). However, it has been suggested that members of this group may be metabolically mixotrophic with the ability to use also organic carbon. Alternatively, they may be using organosulfur compounds such as dimethyl sulphide (DMS), an important product of marine algae (Marshall and Morris, 2012), as electron donors. This metabolic plasticity may contribute to broaden their ecological distribution within the water column, depending on their dominant metabolism.

### Verrucomicrobia

Members of the Verrucomicrobia have recently been shown to degrade polysaccharides and may play an important role in the carbon cycle of freshwater, marine and soil environments (Cardman *et al.*, 2014). Most of the isolated and reconstructed genomes of marine Verrucomicrobia came from surface samples (Yoon *et al.*, 2007; Mavromatis *et al.*, 2010; Hugerth *et al.*, 2015). For the phylogenomic classification of the bins we have included representatives of reconstructed genomes from the Baltic Sea (Hugerth *et al.*, 2015), single cell genomes, and genomes from the family Opitutaceae. The resulting phylogenomic tree (Supplementary Figure 9) showed that our three bins cluster with a reconstructed genome from the Baltic Sea (Opitutaceae bacterium BACL24 MAG 120322 bin51) and the reference genome *Coraliomargarita akajimensis* DSM 45211 (family Puniceicoccaceae), albeit with low ANI values (*ca*. 75%). MED-G30 GC content was similar to *C. akajimensis* (*ca*. 53%) while MED-G31 and MED-G32 showed lower GC values (42.9 and 38.3, respectively). They were all localized in the UP (Figure 2a). These genomes encoded a large number of peptidases, esterases, glycoside hydrolases, carbohydrate lyases and sulfatases (e.g. iduronate-2-sulfatase, arylsulfatase A, arylsulfatase B), providing these microbes with the capacity of degrading complex carbohydrates, as shown for other marine Verrucomicrobia (Cardman *et al.*, 2014). Sulfatases hydrolyze sulfate esters and may be important to utilize sulphated polysaccharides very prevalent in marine algae (Painter, 1983). Interestingly, the presence of the *glg*ABP genes involved in glycogen synthesis in our genomes indicate that they can store glucose in the form of glycogen, as described previously for a verrucomicrobial methanotroph isolated from a microbial mat (Khadem *et al.*, 2012) and in a hypersaline lake (Spring *et al.*, 2016).

### Archaea

Six reconstructed genomes belonged to the marine Euryarchaeota. Further analysis confirmed that all of them belonged to the same cluster (Supplementary Figure 10), marine group II Euryarchaeota, known to be prevalent throughout the photic zone (Martin-Cuadrado *et al.*, 2014). Whereas MED-G33 was closely related to MG2-GG3 (Iverson *et al.*, 2012), assembled from an estuary in the North Pacific, the remaining bins MED-G34 to MED-G38 were related with the recently described Thalassoarchaea (Martin-Cuadrado *et al.*, 2014) assembled from the western Mediterranean Sea, near our sampling site. None of the reconstructed genomes recruited from 1000 m (Figure 2) but most appeared at the lower photic zone (DCM and LP) and only MED-G38 was present at 30 and 60m. At 45m only the previously assembled Thalassoarchaea bin recruited with RPKG values >40, showing a remarkable preference for that depth. Although we could not assign any contig to the marine group III Euryarchaeota, the recruitment of the previously described EUIII-Epi2 and EUIII-Epi6 (Haro-Moreno *et al.*, 2017) showed that they were also present at the DCM. *Ca*. Nitrosopelagicus brevis (Santoro *et al.*, 2015) was the only thaumarchaeon that recruited significantly in our metagenomes and appeared to be restricted to the LP (Figure 2a). The presence of these presumably ammonia-oxidizing archaea was correlated with the detection of oxidized nitrogen compounds (nitrite and nitrate) and with a dramatic reduction of available ammonium (Table 1). Although the affinity for ammonium of some members of this clade may allow them to compete with phototrophs (Martens-Habbena *et al.*, 2009), it is unclear whether all members of the Thaumarchaeota share this characteristic. Furthermore, ammonia-oxidizing archaea have been shown to be inhibited by light (Merbt *et al.*, 2012; French *et al.*, 2012), which may also explain their absence from the upper reaches of the photic zone.

### Depth stratification of rhodopsins

Rhodopsins have been shown to be among the most widespread genes in the photic zone worldwide (Fuhrman *et al.*, 2008; Pinhassi *et al.*, 2016). They are very diverse and distributed throughout most taxa. We evaluated the numbers of rhodopsins among the individual reads and calculated their frequency per genome, normalizing by the number of single copy housekeeping genes (*rec*A and *rad*A) found in the sample (Figure 4b). The total numbers of rhodopsin-assigned reads were clearly correlated to light intensity, from a maximum at 15 m where *ca*. 45% of the genomes contain a rhodopsin, and decreasing with depth.

**Figure 4.**
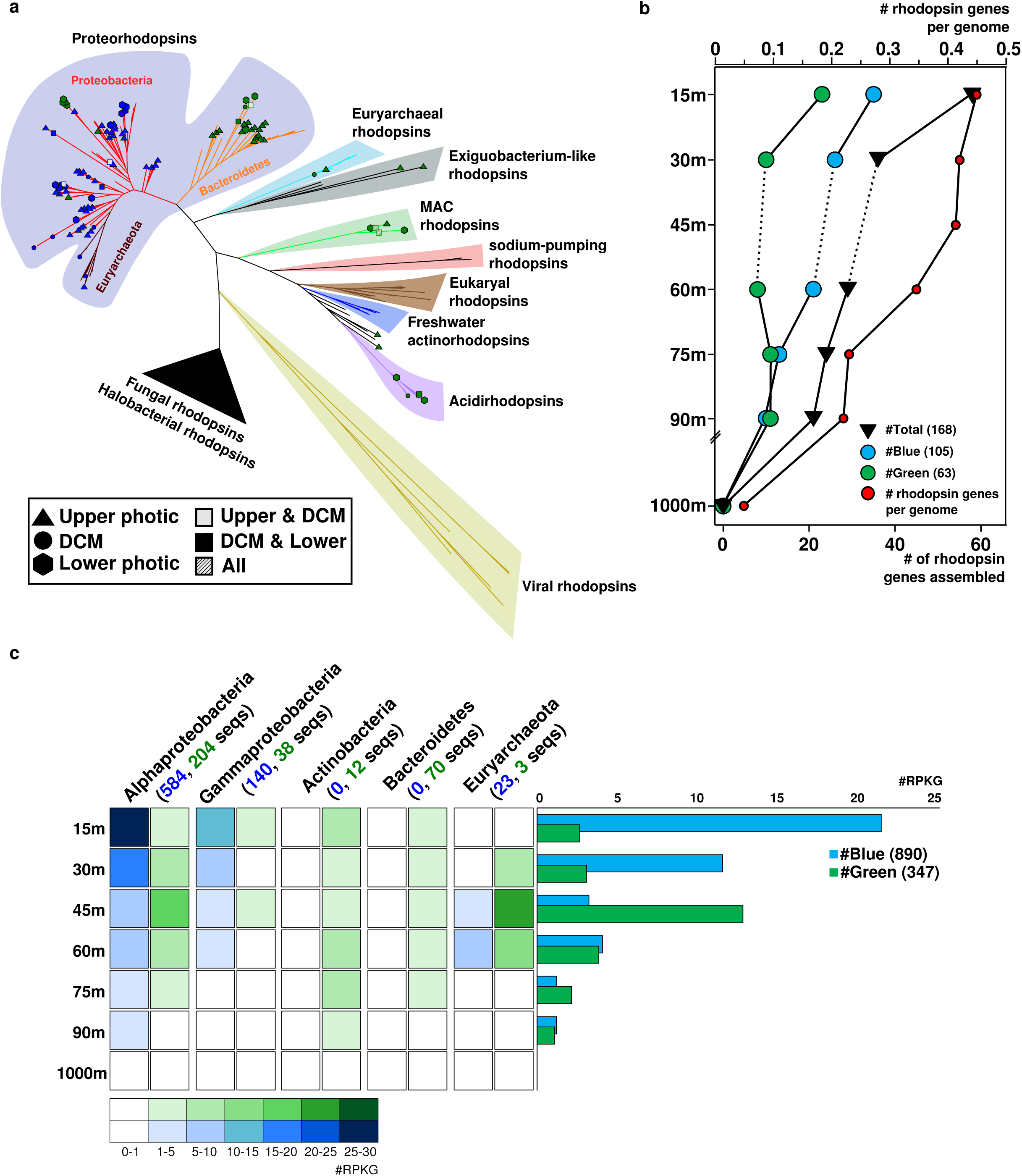
a) Phylogenetic tree of rhodopsin genes detected in the different depth metagenomes. Sequences obtained in this work are represented according to depth and colour absorption (green or blue). b) Number of assembled rhodopsin genes at different depths. Reads annotated as rhodopsin normalized by the number of *rec*A+*rad*A (estimated rhodopsin genes per genome) are indicated by red circles. c) Recruitment of all rhodopsins including those obtained in this work together with the publicly available in the MicRhoDE database. Left panel, rhodopsins classified according to their taxa of origin. Right panel, average number of blue/green rhodopsins for each depth.

We assembled 168 rhodopsin genes throughout the water column. All were classified at least to the phylum level based on flanking genes (Figure 4a and b). Phylogenetic analysis revealed the enormous diversity of this gene family and at least eleven major evolutionary lineages. All the assembled rhodopsin genes clustered with previously described groups indicating that surveys may have be saturating the extant diversity of rhodopsins, at least in the oligotrophic ocean photic zone. Rhodopsin sequences clustered primarily by phylum, with the exception of euryarchaeal rhodopsins as previously reported (Iverson *et al.*, 2012; Haro-Moreno *et al.*, 2017). Within the proteorhodopsin cluster, we could clearly differentiate another bacterial cluster including only Bacteroidetes sequences (Figure 4a). Within clusters, rhodopsin sequences also clustered by depth, with many branches containing only upper or lower photic zone varieties. This confirms the stenobathic character of most groups at the finer level of diversity resolution.

Rhodopsin genes from our metagenomic assemblies and from the MicRhoDE database (Boeuf *et al.*, 2015) were used to recruit reads from the different depths (Figure 4c). We saw no correlation between the predicted absorption spectrum (blue versus green light) of the rhodopsins and the depth from which they recruited the most reads. In contrast, we did see a consistent pattern correlation between the absorption spectrum and the phylogenetic affiliation of the host genome: Bacteroidetes and Actinobacteria all carry green rhodopsins while Proteobacteria have largely the blue variety. The findings suggest, as previously reported (Sabehi *et al.*, 2007; Pinhassi *et al.*, 2016), that the spectral tuning of rhodopsins may not be related to depth adaptation but may involve other physiological characteristics that remain undiscovered.

## DISCUSSION

Stratified systems are widespread on Earth, from microbial mats to meromictic lakes and the temperate ocean. All have in common strong vertical gradients of physico-chemical parameters (Christaki *et al.*, 2011; Bolhuis *et al.*, 2014). The enormous scale of the oceanic environment make the upper 100 m seem relatively small but throughout this stretch of the water column is where the most significant changes occur (Scalan and West, 2002). In contrast, the aphotic zone is much more homogenous and, in the case of the Mediterranean, which has no permanent thermocline (D’Ortenzio *et al.*, 2005), likely even more so. The main variation would be the decrease in DOM due to the gradual consumption by the microbiota that is not regenerated by primary productivity.

The vast majority of the microbial genomes recruited very unevenly, particularly considering that all the samples were collected on the same day (except the 1000m). All the genomes recruited much more at one single specific depth and most (*ca*. 70%) recruited only at either one or two contiguous depth metagenomes. This indicates that the distribution of most of these microbes extends only over a 30 m thick layer within the *ca*. 100 m deep photic zone. Only one of the photic zone genomes, Sphingomonadaceae MED-G03, recruited at the 1000m sample. This genome recruited actually more at this depth and it could be the only truly eurybathic microbe among the ones assembled here. The actinobacterial genomes seemed to be the next most eurybathic and, although they were always more prevalent at a single depth, they were detectable at four depths, with the exception of the single cell genome SCGC-AAA015-M09 (Swan *et al.*, 2013) (only found at 15 and 30m). The Alphaproteobacteria (with the exception of the *Pelagibacterales*), like most *Bacteroidetes* and Gammaproteobacteria were only detected at one or two depths. Of particular note, Pelagibacter MED-G39 was found exclusively in the Mediterranean Sea, a truly endemic microorganism, and was also the most abundant genome in samples throughout this body of water.

We also found biological stratification at multiple levels of diversity, from higher taxonomic levels (class, orders and families), to closely related microbes belonging probably to the same species (Figures 1 and 2). In addition to these layers of diversity, there are still others associated with the flexible genomes (Rodriguez-Valera *et al.*, 2016) that cannot be assembled from metagenomics datasets (because it varies from one clone to another) that allow each taxonomic unit of the species rank to adjust to microniche diversity (Kashtan *et al.*, 2014; Rodriguez-Valera *et al.*, 2009). Our work illustrates again the enormous diversity displayed by prokaryotic cells in an environment considered relatively homogeneous.

It is worthwhile noting that the Mediterranean only remains stratified during 6-7 months and that the water column is nearly fully mixed the rest of the year. Therefore, it is likely that the fine stratification discovered for many of these microbes develops over a relatively short time span. Deep dwellers such as the Euryarchaea of MGII (Hugoni *et al.*, 2013; Martin-Cuadrado *et al.*, 2014) that are transported to the surface in winter, disappear or migrate during the summer to be found only in the LP, whereas the opposite happens to surface dwellers. This finding has important ramifications for global marine studies that most often take only surface or, at most, one single subsurface photic zone sample. Furthermore, we believe that surface samples should be avoided whenever possible, particularly in stratified water columns, since they tend to be extremely patchy and are also potentially contaminated by the proximity of the research vessel/sampling device. What is more, most microbes are likely less active at the surface, at least during the summer, due to the strong UV light and the depletion of nutrients (Moore *et al.*, 2002; Moutin and Raimbault, 2002). Depth in the water column is, as shown here, the single most important factor that structures the community in the open ocean (Delong *et al.*, 2006). Therefore, fine profiles like the one described here are essential before other likely less critical gradients are approached.

## Acknowledgements.

This work was supported by projects MEDIMAX BFPU2013-48007-P, VIREVO CGL2016-76273-P and Acciones de dinaminación REDES DE EXCELENCIA CONSOLIDER CGL2015- 71523-REDC, from the Spanish Ministerio de Economía y Competitividad; and project AQUAMET PROMETEOII/2014/012 from Generalitat Valenciana. JHM was supported with a PhD fellowship from the Spanish Ministerio de Economía y Competitividad. MLP was supported with a Postdoctoral fellowship from the Valencian Consellería de Educació, Investigació, Cultura i Esport (APOSTD/2016/051).

## Author Contributions

FRV conceived the study, helped with the analysis and wrote most of the manuscript. JHM analysed together with MLP the data. AC and AP helped in the sampling and analyzed all physicochemical and ecological parameters. JRT helped to analyze the data and write the manuscript.

## References

Albertsen M, Hugenholtz P, Skarshewski A, Nielsen KL, Tyson GW, Nielsen PH. (2013). Genome sequences of rare, uncultured bacteria obtained by differential coverage binning of multiple metagenomes. Nat Biotechnol 31: 533–8.

Altschul SF, Madden TL, Schäffer AA, Zhang J, Zhang Z, Miller W, et al. (1997). Gapped BLAST and PSI-BLAST: A new generation of protein database search programs. Nucleic Acids Res 25: 3389–3402.

Balkwill D, Fredrickson JK, Romine MF. (2006). Sphingomonas and Related Genera. Prokaryotes SE - 23 605–629.

Bankevich A, Nurk S, Antipov D, Gurevich AA, Dvorkin M, Kulikov AS, et al. (2012). SPAdes: A New Genome Assembly Algorithm and Its Applications to Single-Cell Sequencing. J Comput Biol 19: 455–477.

Bendall ML, Stevens SL, Chan L-K, Malfatti S, Schwientek P, Tremblay J, et al. (2016). Genome-wide selective sweeps and gene-specific sweeps in natural bacterial populations. ISME J 10: 1–13.

Biller SJ, Berube PM, Lindell D, Chisholm SW. (2014). Prochlorococcus: the structure and function of collective diversity. Nat Rev Microbiol 13: 13–27.

Boeuf D, Audic S, Brillet-Guéguen L, Caron C, Jeanthon C. (2015). MicRhoDE: A curated database for the analysis of microbial rhodopsin diversity and evolution. Database 2015. e-pub ahead of print, doi:10.1093/database/bav080.

Bolhuis H, Cretoiu MS, Stal LJ. (2014). Molecular ecology of microbial mats. FEMS Microbiol Ecol 90: 335–350.

Buchfink B, Xie C, Huson DH. (2015). Fast and sensitive protein alignment using DIAMOND. Nat Methods 12: 59–60.

Cardman Z, Arnosti C, Durbin A, Ziervogel K, Cox C, Steen AD, et al. (2014). Verrucomicrobia are candidates for polysaccharide-degrading bacterioplankton in an Arctic fjord of Svalbard. Appl Environ Microbiol 80: 3749–3756.

Cho JC, Giovannoni SJ. (2004). Cultivation and Growth Characteristics of a Diverse Group of Oligotrophic Marine Gammaproteobacteria. Appl Environ Microbiol 70: 432–440.

Christaki U, Van Wambeke F, Lefevre D, Lagaria A, Prieur L, Pujo-Pay M, et al. (2011). Microbial food webs and metabolic state across oligotrophic waters of the Mediterranean Sea during summer. Biogeosciences 8: 1839–1852.

Cole JR, Wang Q, Fish JA, Chai B, McGarrell DM, Sun Y, et al. (2014). Ribosomal Database Project: Data and tools for high throughput rRNA analysis. Nucleic Acids Res 42: 633–642.

Cram J a, Chow C-ET, Sachdeva R, Needham DM, Parada AE, Steele J a, et al. (2014). Seasonal and interannual variability of the marine bacterioplankton community throughout the water column over ten years. ISME J 1–18.

D’Ortenzio F, Iudicone D, de Boyer Montegut C, Testor P, Antoine D, Marullo S, et al. (2005). Seasonal variability of the mixed layer depth in the Mediterranean Sea as derived from in situ profiles. Geophys Res Lett 32: 1–4.

Delong EF, Preston CM, Mincer T, Rich V, Hallam SJ, Frigaard N, et al. (2006). Community Genomics among microbial assemblages in the Ocean’ s Interior. Science (80-) 311: 496–503.

Denaro G, Valenti D, La Cognata A, Spagnolo B, Bonanno A, Basilone G, et al. (2013). Spatio-temporal behaviour of the deep chlorophyll maximum in Mediterranean Sea: Development of a stochastic model for picophytoplankton dynamics. Ecol Complex 13: 21–34.

Eaton AD, Clesceri LS, Greenberg AE, Franson MAH, Association APH, Association AWW, et al. (1992). Standard Methods For the Examination of Water and Wastewater.

Eddy SR. (1995). Multiple alignment using hidden Markov models. Proc Int Conf Intell Syst Mol Biol 3: 114–120.

Edgar RC. (2004). MUSCLE: Multiple sequence alignment with high accuracy and high throughput. Nucleic Acids Res 32: 1792–1797.

Edgar RC. (2010). Search and clustering orders of magnitude faster than BLAST. Bioinformatics 26: 2460–2461.

Estrada M, Henriksen P, Gasol JM, Casamayor EO, Pedrós-Alió C. (2004). Diversity of planktonic photoautotrophic microorganisms along a salinity gradient as depicted by microscopy, flow cytometry, pigment analysis and DNA-based methods. FEMS Microbiol Ecol 49: 281–293.

Fernández-Gómez B, Richter M, Schüler M, Pinhassi J, Acinas SG, González JM, et al. (2013). Ecology of marine Bacteroidetes: a comparative genomics approach. ISME J 7: 1026–37.

Ferreira AJS, Siam R, Setubal JC, Moustafa A, Sayed A, Chambergo FS, et al. (2014). Core microbial functional activities in ocean environments revealed by global metagenomic profiling analyses. PLoS One 9. e-pub ahead of print, doi:10.1371/journal.pone.0097338.

Field CB, Behrenfeld MJ, Randerson JT, Falkowski P. (1998). Primary production of the biosphere: integrating terrestrial and oceanic components. Science 281: 237–40.

French E, Kozlowski JA, Mukherjee M, Bullerjahn G, Bollmann A. (2012). Ecophysiological characterization of ammonia-oxidizing archaea and bacteria from freshwater. Appl Environ Microbiol 78: 5773–5780.

Fuhrman J a, Schwalbach MS, Stingl U. (2008). Proteorhodopsins: an array of physiological roles? Nat Rev Microbiol 6: 488–494.

Garczarek L, Dufresne A, Rousvoal S, West NJ, Mazard S, Marie D, et al. (2007). High vertical and low horizontal diversity of Prochlorococcus ecotypes in the Mediterranean Sea in summer. FEMS Microbiol Ecol 60: 189–206.

Ghai R, Martin-Cuadrado A-B, Molto AG, Heredia IG, Cabrera R, Martin J, et al. (2010). Metagenome of the Mediterranean deep chlorophyll maximum studied by direct and fosmid library 454 pyrosequencing. ISME J 4: 1154–1166.

Ghai R, Mizuno CM, Picazo A, Camacho A, Rodriguez-Valera F. (2013). Metagenomics uncovers a new group of low GC and ultra-small marine Actinobacteria. Sci Rep 3: 2471.

Giovannoni SJ, Vergin KL. (2012). Seasonality in Ocean Microbial Communities. Science (80-) 335: 671–676.

Glaubitz S, Kießlich K, Meeske C, Labrenz M, Jürgens K. (2013). SUP05 Dominates the gammaproteobacterial sulfur oxidizer assemblages in pelagic redoxclines of the central baltic and black seas. Appl Environ Microbiol 79: 2767–2776.

Gonzaga A, Martin-Cuadrado AB, López-Pérez M, Mizuno CM, García-Heredia I, Kimes NE, et al. (2012). Polyclonality of concurrent natural populations of Alteromonas macleodii. Genome Biol Evol 4: 1360–1374.

Grote J, Thrash JC, Huggett MJ. (2012). Streamlining and Core Genome Conservation among Highly Divergent Members of the SAR11 Clade. 3: 1–13.

Haft DH, Loftus BJ, Richardson DL, Yang F, Eisen JA, Paulsen IT, et al. (2001). TIGRFAMs: a protein family resource for the functional identification of proteins. Nucleic Acids Res 29: 41–43.

Harmelin JG, Dhondt JL. (1993). Transfers of Bryozoan Species between the Atlantic-Ocean and the Mediterranean-Sea Via the Strait of Gibraltar. Oceanol Acta 16: 63–72.

Haro-Moreno JM, Rodriguez-Valera F, López-García P, Moreira D, Martin-Cuadrado A-B. (2017). New insights into marine group III Euryarchaeota, from dark to light. ISME J 1–16.

Huang Y, Gilna P, Li W. (2009). Identification of ribosomal RNA genes in metagenomic fragments. Bioinformatics 25: 1338–1340.

Hugerth LW, Larsson J, Alneberg J, Lindh M V, Legrand C, Pinhassi J, et al. (2015). Metagenome-assembled genomes uncover a global brackish microbiome. bioRxiv 18465.

Hugoni M, Taib N, Debroas D, Domaizon I, Jouan Dufournel I, Bronner G, et al. (2013). Structure of the rare archaeal biosphere and seasonal dynamics of active ecotypes in surface coastal waters. Proc Natl Acad Sci U S A 110: 6004–9.

Huson DH, Beier S, Flade I, Grska A, El-Hadidi M, Mitra S, et al. (2016). MEGAN Community Edition - Interactive Exploration and Analysis of Large-Scale Microbiome Sequencing Data. PLoS Comput Biol 12. e-pub ahead of print, doi:10.1371/journal.pcbi.1004957.

Hyatt D, Chen G-L, Locascio PF, Land ML, Larimer FW, Hauser LJ. (2010). Prodigal: prokaryotic gene recognition and translation initiation site identification. BMC Bioinformatics 11: 119.

Iverson V, Morris RM, Frazar CD, Berthiaume CT, Morales RL, Armbrust E V. (2012). Untangling Genomes from Metagenomes: Revealing an Uncultured Class of Marine Euryarchaeota. Science (80-) 335: 587–590.

Jiao N, Zhang F, Hong N. (2010). Significant roles of bacteriochlorophylla supplemental to chlorophylla in the ocean. ISME J 4: 595–597.

Jimenez-Infante F, Ngugi DK, Alam I, Rashid M, Baalawi W, Kamau A a., et al. (2014). Genomic differentiation among two strains of the PS1 clade isolated from geographically separated marine habitats. FEMS Microbiol Ecol 89: 181–197.

Kashtan N, Roggensack SE, Rodrigue S, Thompson JW, Biller SJ, Coe A, et al. (2014). Single-cell genomics reveals hundreds of coexisting subpopulations in wild Prochlorococcus. Science 344: 416–20.

Khadem AF, Van Teeseling MCF, Van Niftrik L, Jetten MSM, Op Den Camp HJM, Pol A. (2012). Genomic and physiological analysis of carbon storage in the verrucomicrobial methanotroph ‘Ca. Methylacidiphilum fumariolicum’ SolV. Front Microbiol 3. e-pub ahead of print, doi:10.3389/fmicb.2012.00345.

Konstantinidis KT, Braff J, Karl DM, DeLong EF. (2009). Comparative metagenomic analysis of a microbial community residing at a depth of 4,000 meters at station ALOHA in the North Pacific Subtropical Gyre. Appl Environ Microbiol 75: 5345–5355.

Kumar S, Stecher G, Tamura K. (2016). MEGA7: Molecular Evolutionary Genetics Analysis version 7.0 for bigger datasets. Mol Biol Evol 33: msw054.

Lassmann T, Sonnhammer ELL. (2005). Kalign--an accurate and fast multiple sequence alignment algorithm. BMC Bioinformatics 6: 298.

Lê S, Josse J, Husson F. (2008). FactoMineR: An R Package for Multivariate Analysis. J Stat Softw 25: 1–18.

Letelier RM, Karl DM, Abbott MR, Bidigare RR. (2004). Light driven seasonal patterns of chlorophyll and nitrate in the lower euphotic zone of the North Pacific Subtropical Gyre. Limnol Oceanogr 49: 508–519.

Li H, Durbin R. (2009). Fast and accurate short read alignment with Burrows-Wheeler transform. Bioinformatics 25: 1754–1760.

López-Pérez M, Kimes NE, Haro-Moreno JM, Rodriguez-valera F. (2016). Not All Particles Are Equal?: The Selective Enrichment of Particle-Associated Bacteria from the Mediterranean Sea. 7. e-pub ahead of print, doi:10.3389/fmicb.2016.00996.

Lowe TM, Eddy SR. (1996). TRNAscan-SE: A program for improved detection of transfer RNA genes in genomic sequence. Nucleic Acids Res 25: 955–964.

Luo H, Thompson LR, Stingl U, Hughes AL. (2015). Selection maintains low genomic GC content in marine SAR11 lineages. Mol Biol Evol 32: 2738–2748.

Man D, Wang W, Sabehi G, Aravind L, Post AF, Massana R, et al. (2003). Diversification and spectral tuning in marine proteorhodopsins. EMBO J 22: 1725–1731.

Marie D, Partensky F, Jacquet S, Vaulot D. (1997). Enumeration and cell cycle analysis of natural populations of marine picoplankton by flow cytometry using the nucleic acid stain SYBR Green I. Appl Environ Microbiol 63: 186–193.

Marshall KT, Morris RM. (2012). Isolation of an aerobic sulfur oxidizer from the SUP05/Arctic96BD-19 clade. ISME J 7: 452–455.

Martens-Habbena W, Berube PM, Urakawa H, de la Torre JR, Stahl DA, Torre J, et al. (2009). Ammonia oxidation kinetics determine niche separation of nitrifying Archaea and Bacteria. Nature 461: 976–979.

Martin-Cuadrado A-B, Garcia-Heredia I, Moltó AG, López-Úbeda R, Kimes N, López-García P, et al. (2014). A new class of marine Euryarchaeota group II from the mediterranean deep chlorophyll maximum. ISME J 1–16.

Martin-Cuadrado A-B, Rodriguez-Valera F, Moreira D, Alba JC, Ivars-Martínez E, Henn MR, et al. (2008). Hindsight in the relative abundance, metabolic potential and genome dynamics of uncultivated marine archaea from comparative metagenomic analyses of bathypelagic plankton of different oceanic regions. ISME J 2: 865–886.

Mavromatis K, Abt B, Brambilla E, Lapidus A, Copeland A, Deshpande S, et al. (2010). Complete genome sequence of Coraliomargarita akajimensis type strain (04OKA010-24). Stand Genomic Sci 2: 290–299.

Mella-Flores D, Mazard S, Humily F, Partensky F, Mahé F, Bariat L, et al. (2011). Is the distribution of Prochlorococcus and Synechococcus ecotypes in the Mediterranean Sea affected by global warming? Biogeosciences 8: 2785–2804.

Merbt SN, Stahl DA, Casamayor EO, Martfi E, Nicol GW, Prosser JI. (2012). Differential photoinhibition of bacterial and archaeal ammonia oxidation. FEMS Microbiol Lett 327: 41–46.

Mizuno CM, Rodriguez-valera F, Ghai R. (2015). Genomes of Planktonic Acidimicrobiales?: Widening Horizons for Marine Actinobacteria by Metagenomics. 6: 1–11.

Moore JK, Doney SC, Glover DM, Fung IY. (2002). Iron cycling and nutrient-limitation patterns in surface waters of the world ocean. Deep Res Part II Top Stud Oceanogr 49: 463–507.

Moutin T, Raimbault P. (2002). Primary production, carbon export and nutrients availability in western and eastern Mediterranean Sea in early summer 1996 (MINOS cruise). J Mar Syst 33-34: 273–288.

Munoz R, Rosselló-Móra R, Amann R. (2016). Revised phylogeny of Bacteroidetes and proposal of sixteen new taxa and two new combinations including Rhodothermaeota phyl. nov. Syst Appl Microbiol 39: 281–296.

Nawrocki EP. (2009). Structural RNA Homology Search and Alignment Using Covariance Models. 281.

Nayfach S, Pollard KS. (2015). Average genome size estimation improves comparative metagenomics and sheds light on the functional ecology of the human microbiome. Genome Biol 16: 51.

Painter TJ. (1983). 4 - Algal Polysaccharides. In: The Polysaccharides. pp 195–285.

Parks DH, Imelfort M, Skennerton CT, Hugenholtz P, Tyson GW. (2015). CheckM: assessing the quality of microbial genomes recovered from isolates, single cells, and metagenomes. Genome Res 25: 1043–55.

Partensky F, Blanchot J, Vaulot D. (1999). Differential distribution and ecology of Prochlorococcus and Synechococcus in oceanic waters?: a review. Bull l’Institut océanographique 19: 457–475.

Peng Y, Leung HCM, Yiu SM, Chin FYL. (2012). IDBA-UD: A de novo assembler for single-cell and metagenomic sequencing data with highly uneven depth. Bioinformatics 28: 1420–1428.

Picazo A, Rochera C, Vicente E, Miracle MR, Camacho A. (2013). Spectrophotometric methods for the determination of photosynthetic pigments in stratified lakes: A critical analysis based on comparisons with HPLC determinations in a model lake. Limnetica 32: 139–158.

Pinhassi J, DeLong EF, Béjà O, González JM, Pedrós-Alió C. (2016). Marine Bacterial and Archaeal Ion-Pumping Rhodopsins: Genetic Diversity, Physiology, and Ecology. Microbiol Mol Biol Rev 80: 929–54.

Raes J, Korbel JO, Lercher MJ, von Mering C, Bork P. (2007). Prediction of effective genome size in metagenomic samples. Genome Biol 8: R10.

Rice P, Longden I, Bleasby A. (2000). EMBOSS: The European Molecular Biology Open Software Suite. Trends Genet 16: 276–277.

Robert C Edgar. (2010). UCLUST: Extreme high-speed sequence clustering, alignment and database search. http://www.drive5.com/usearch.

Rodriguez-r LM, Konstantinidis KT. (2016). The enveomics collection?: a toolbox for specialized analyses of microbial genomes and metagenomes. Peer J Prepr. e-pub ahead of print, doi:10.7287/peerj.preprints.1900v1.

Rodriguez-Valera F, Martin-Cuadrado A-B, López-Pérez M. (2016). Flexible genomic islands as drivers of genome evolution. Curr Opin Microbiol 31: 154–160.

Rodriguez-Valera F, Martin-Cuadrado A-B, Rodriguez-Brito B, Pasić L, Thingstad TF, Rohwer F, et al. (2009). Explaining microbial population genomics through phage predation. Nat Rev Microbiol 7: 828–36.

Rusch DB, Halpern AL, Sutton G, Heidelberg KB, Williamson S, Yooseph S, et al. (2007). The Sorcerer II Global Ocean Sampling expedition: Northwest Atlantic through eastern tropical Pacific. PLoS Biol 5: 0398–0431.

Sabehi G, Kirkup BC, Rozenberg M, Stambler N, Polz MF, Béjà O. (2007). Adaptation and spectral tuning in divergent marine proteorhodopsins from the eastern Mediterranean and the Sargasso Seas. Isme J 1: 48–55.

Santoro AE, Dupont CL, Richter RA, Craig MT, Carini P, McIlvin MR, et al. (2015). Genomic and proteomic characterization of ‘Candidatus Nitrosopelagicus brevis’: An ammonia-oxidizing archaeon from the open ocean. Proc Natl Acad Sci 112: 1173–1178.

Scalan DJ, West NJ. (2002). Molecular ecology of the marine cyanobacteria genera Prochlorococcus and Synechococcus. FEMS Microbiol Ecol 40: 1–12.

Shi Y, Tyson GW, Eppley JM, DeLong EF. (2011). Integrated metatranscriptomic and metagenomic analyses of stratified microbial assemblages in the open ocean. ISME J 5: 999–1013.

Spring S, Bunk B, Spröer C, Schumann P, Rohde M, Tindall BJ, et al. (2016). Characterization of the first cultured representative of Verrucomicrobia subdivision 5 indicates the proposal of a novel phylum. ISME J 10: 1–16.

Sunagawa S, Coelho LP, Chaffron S, Kultima JR, Labadie K, Salazar G, et al. (2015). Structure and function of the global ocean microbiome. Science (80-) 348: 1261359.

Swan BK, Martinez-Garcia M, Preston CM, Sczyrba A, Woyke T, Lamy D, et al. (2011). Potential for chemolithoautotrophy among ubiquitous bacteria lineages in the dark ocean. Science 333: 1296–300.

Swan BK, Tupper B, Sczyrba A, Lauro FM, Martinez-Garcia M, González JM, et al. (2013). Prevalent genome streamlining and latitudinal divergence of planktonic bacteria in the surface ocean. Proc Natl Acad Sci U S A 110: 11463–8.

Tatusov RL, Natale DA, Garkavtsev I V, Tatusova TA, Shankavaram UT, Rao BS, et al. (2001). The COG database: new developments in phylogenetic classification of proteins from complete genomes. Nucleic Acids Res 29: 22–8.

Thompson AW, Huang K, Saito MA, Chisholm SW. (2011). Transcriptome response of high and low-light-adapted Prochlorococcus strains to changing iron availability. ISME J 5: 1580–94.

Thrash JC, Temperton B, Swan BK, Landry ZC, Woyke T, DeLong EF, et al. (2014). Single-cell enabled comparative genomics of a deep ocean SAR11 bathytype. ISME J 8: 1440–51.

Tolonen AC, Aach J, Lindell D, Johnson ZI, Rector T, Steen R, et al. (2006). Global gene expression of Prochlorococcus ecotypes in response to changes in nitrogen availability. Mol Syst Biol 2: 53.

Tsementzi D, Wu J, Deutsch S, Nath S, Rodriguez-R LM, Burns AS, et al. (2016). SAR11 bacteria linked to ocean anoxia and nitrogen loss. Nature 536: 179–183.

Viklund J, Martijn J, Ettema TJG, Andersson SGE. (2013). Comparative and phylogenomic evidence that the alphaproteobacterium HIMB59 is not a member of the oceanic SAR11 clade. PLoS One 8. e-pub ahead of print, doi:10.1371/journal.pone.0078858.

Walsh DA, Zaikova E, Howes CG, Song YC, Wright JJ, Tringe SG, et al. (2009). Metagenome of a versatile chemolithoautotroph from expanding oceanic dead zones. Science 326: 578–582.

Walsh EA, Kirkpatrick JB, Rutherford SD, Smith DC, Sogin M, D’Hondt S. (2015). Bacterial diversity and community composition from seasurface to subseafloor. ISME J 10: 1–11.

Yan S, Fuchs BM, Lenk S, Harder J, Wulf J, Jiao NZ, et al. (2009). Biogeography and phylogeny of the NOR5/OM60 clade of Gammaproteobacteria. Syst Appl Microbiol 32: 124–139.

Yoon J, Yasumoto-Hirose M, Katsuta A, Sekiguchi H, Matsuda S, Kasai H, et al. (2007). Coraliomargarita akajimensis gen. nov., sp. nov., a novel member of the phylum ‘Verrucomicrobia’ isolated from seawater in Japan. Int J Syst Evol Microbiol 57: 959–963.

Zwirglmaier K, Heywood JL, Chamberlain K, Woodward EMS, Zubkov M V., Scanlan DJ. (2007). Basin-scale distribution patterns of picocyanobacterial lineages in the Atlantic Ocean. Environ Microbiol 9: 1278–1290.

